# Cellular properties of intrinsically photosensitive retinal ganglion cells during postnatal development

**DOI:** 10.1101/619387

**Authors:** Jasmine A. Lucas, Tiffany M. Schmidt

## Abstract

**Background:** Melanopsin-expressing, intrinsically photosensitive retinal ganglion cells (ipRGCs) respond directly to light and have been shown to mediate a broad variety of visual behaviors in adult animals. ipRGCs are also the first light sensitive cells in the developing retina, and have been implicated in a number of retinal developmental processes such as pruning of retinal vasculature and refinement of retinofugal projections. However, little is currently known about the properties of the six ipRGC subtypes during development, and how these cells act to influence retinal development. We therefore sought to characterize the structure, physiology, and birthdate of the most abundant ipRGC subtypes, M1, M2, and M4, at discrete postnatal developmental timepoints.

**Methods:** We utilized whole cell patch clamp to measure the electrophysiological and morphological properties of ipRGC subtypes through postnatal development. We also used EdU labeling to determine the embryonic timepoints at which ipRGC subtypes terminally differentiate.

**Results:** Our data show that ipRGC subtypes are distinguishable from each other early in postnatal development. Additionally, we find that while ipRGC subtypes terminally differentiate at similar embryonic stages, the subtypes reach adult-like morphology and physiology at different developmental timepoints.

**Conclusions:** This work provides a broad assessment of ipRGC morphological and physiological properties during the postnatal stages at which they are most influential in modulating retinal development, and lays the groundwork for further understanding of the specific role of each ipRGC subtype in influencing retinal and visual system development.

## Background

Melanopsin-expressing, intrinsically photosensitive retinal ganglion cells (ipRGCs) represent a class of non-canonical, ganglion cell photoreceptors. These cells influence a variety of visual behaviors including contrast sensitivity (1), circadian photoentrainment (2–4), sleep (5, 6), and even mood (7, 8). These wide-ranging behavioral influences are attributed to the multiple subtypes (M1-6) that comprise the ipRGC population, with different subtypes possessing a unique complement of cellular properties and playing distinct roles in vision. For example, the M1 ipRGC subtype has been linked to subconscious, non-image forming behaviors including circadian photoentrainment, the pupillary light reflex, and even regulation of mood and learning. The M4 ipRGCs, in comparison, are important for proper contrast sensitivity in visual perception (1, 9).

Although ipRGCs have been categorized based on their adult characteristics, they are in fact light sensitive from embryonic stages (10–12) and begin to exhibit diverse light response properties at early postnatal stages (13, 14). Thus, these unique photoreceptors are light sensitive long before the rest of the retinal circuitry is able to functionally relay rod/cone signals around ~P12 when the eyes open (15, 16). This early photosensitivity has led to multiple studies examining potential developmental influences of ipRGCs on the developing retina and visual system. One study found that melanopsin modulates the branching patterns of retinal vasculature in a light-dependent manner (10). Other studies revealed that melanopsin and ipRGCs can influence spontaneous retinal waves (17) and that they are important retinofugal refinement (17, 18). Surprisingly, light and melanopsin can even drive a light avoidance behavior in neonatal mice as young as 6 days old (19).

While it is clear that light is modulating retinal development and even pup behavior through melanopsin, the circuit mechanisms of these effects remain unclear. In particular, it is not known which of the six ipRGC subtypes mediate these developmental effects. A first step in determining the role of the ipRGC subtypes in development is characterizing the developmental time course of the maturation of each cell type. A previous study has revealed that there are at least three physiological ipRGC subtypes during development, type I, II, and III (13). These subtypes were differentiated based on the size and sensitivity of their light responses with follow up studies proposing that that the type I corresponds to the adult M4 subtype, type II to the M2 subtype, and type III to the M1 subtype (14, 20). Beyond this, little is known about the structure and function of ipRGC subtypes during development, and yet this information is a necessary first step in understanding the mechanisms by which ipRGCs influence the developing retina. We therefore set out to characterize the morphology, physiology, and developmental “birth” date of the three major ipRGC subtypes, M1, M2, and M4. We found that ipRGC subtypes are differentiable at early postnatal stages and seem to exhibit different rates of maturation. Moreover, we find that while ipRGCs are generally born at similar embryonic time points, their birth is largely complete at timepoints earlier than two groups of conventional RGCs: the OFF alpha RGCs and Brn3a positive RGCs.

## Methods

### Animals

All procedures were approved by the Animal Care and Use Committee at Northwestern University. Both male and female mice were used and are from a mixed B6/129 background. Adult mice were between 30-60 days of age.

### Electrophysiology

We used *Opn4-GFP* (21) mice for all electrophysiological recordings. All mice P14 and under were dark adapted 1-2 hrs prior to recording. Adult mice were dark adapted overnight. Pups aged P10 and under were sacrificed via decapitation. P14 pups and adult mice were euthanized using CO_2_ asphyxiation followed by cervical dislocation under dim red illumination. Eyes were enucleated, and retina were dissected under dim red light in carbogenated (95% O_2_-5%CO_2_) Ame’s medium (Sigma, A1420). Retinas were then sliced in half and incubated at 25°C in Ame’s solution for at least 30min. Retinas were mounted ganglion side up on glass bottom recording chamber and anchored using a platinum ring with nylon mesh. Recordings were performed at 24-26 °C with 1-2mL/min flow of Ame’s solution. ipRGCs (GFP positive) were visualized using whole field 480 nm light for less than 30 seconds at 3.5×10^17^ photons/cm^2^s^−1^ intensity, and so all properties of ipRGCs were measured in light adapted tissue. Adult M4 cells were targeted using their characteristic large somata and confirmed post-recording with immunohistochemistry and dendritic stratification.

Recording pipettes were between 4-8 MΩ and filled with following internal solution (in mM): 125 K-gluconate, 2 CaCl2, 2 MgCl2, 10 EGTA, 10 HEPES, 2 Na2-ATP, 0.5 Na-GTP, 10µM Alexa Fluor hydrazide 488 (Thermo, A10436), and 0.3% neurobiotin (Vector, SP-1120-50), pH to 7.2 with KOH.

After recording, retina pieces were fixed with 4% PFA overnight. Pieces were then washed with PBS, blocked for 1hr in 0.3% Triton-X, 6% donkey serum at room temp. After block then placed in the following primary for 2 nights. On the third day, retina pieces were washed with PBS and placed into the following secondary solution for 2hrs at room temperature in the dark. Retinas were then washed and mounted in fluoromount (Sigma, F4680). See table 1 for specific antibodies and concentrations. All images were captured using a confocal laser scanning microscope (LSM, DFC 310 FX, Leica) with a 40x oil-immersion objective.

### Inner Plexiform Lamination Analysis

Dendritic arbors from ipRGCs were traced using Fiji plugin software, simple neurite tracer with subsequent analysis done by using a similar program and methods as described in Nath & Schwartz, 2016 (22, 23).

### Morphological Analysis

FIJI (ImageJ) software was used to analyze cell morphology. For soma diameter measurements, we took a DIC image of the soma before patching. Using the polygon tool, we traced the entire soma and calculated the diameter using the circle equation. A similar method was used to calculate dendritic diameter from cell fill images. We used FIJI plugin, neuronJ, to trace cell fills to get a measurement of total dendritic length. These traced cell fills were subsequently used for Sholl analysis which was performed using the FIJI software.

### Electrophysiological Analysis

#### C_m_/R_inp_

Cells were given a 10mV hyperpolarization step in voltage clamp mode. Capacitance and input resistance were calculated from recorded trace using Ohm’s law.

#### V_m_

Cells were recorded at rest in current clamp mode for 3 minutes with the last minute of the recording being averaged to yield the resting membrane potential. Spike frequency was also assessed in the last minute of the recording.

#### Depolarizing current injections

Current was injected to hold cells at −79 mV and cells were then injected with 1s of +10pA or +20pA stepwise current until cells reached a current that caused depolarization block.

#### Action Potentials

For action potential analysis, the first action potential elicited at the lowest depolarizing current was used for full width half maximum (from threshold), threshold (24), and hyperpolarization analysis. Hyperpolarization was difference between the threshold and the lowest point following action potential peak.

#### Light Onset

Light onset was defined when cell membrane potential reach 50% of maximum light response during the lights on period.

### Statistics

Using Prism graphpad, we analyzed data with non-parametric one-way (Kruskal-Wallis) with Dunn’s multiple comparisons test for any ANOVA that indicated statistical difference. Statistical significance was concluded when p <0.05.

### Birthdating

We crossed Opn4^Cre/+^;ZEG to Wildtype mice to generate Opn4^Cre/+^;ZEG animals and Opn4^LacZ/LacZ^ to *Opn4-GFP* to generate Opn4^LacZ/+^;*Opn4-GFP* mice. Male and female mice were house together and female mice were checked daily for copulation plug. Once plug was confirmed, the potentially pregnant female was separated from the male and singly housed. On the day before the targeted gestation day, pregnant females were water deprived for 24hrs. On the targeted gestation day, pregnant females were given 400µL of water containing 30µg/g of EdU (Abcam, ab146186) every 2 hrs for 12 hours. Gestation day was confirmed when female gave birth on the 19^th^ day.

EdU mice of the correct genotype were dissected between 30-60 days of age. Mice were euthanized with CO_2_ asphyxiation, followed by cervical dislocation. Eyes were enucleated and retinas were fixed overnight in 4% PFA at 4°C. The next day retinas were washed with PBS, blocked for 1hr at room temperature in 0.3% Triton-X, 6% goat or donkey serum and then placed into primary solution for 2-3 nights at 4°C. Then retinas were washed with PBS and incubated in secondary solution for 2 hours at room temperature. Finally, retinas were washed with PBS and click-it reaction was performed according to manufacturer’s specifications on flat mounted retina (Thermo, C10640). After click-it reaction, retinas were washed with PBS and mounted in fluoromount. See table 1 for specific antibodies and concentrations. 8 images were acquired at 0.5mm, 1.0mm, and 1.5mm from the optic nerve for a total of 24 per retina. All images were captured using a confocal laser scanning microscope (LSM, DFC 310 FX, Leica) with a 40x oil-immersion objective.

**Table 1:** Antibodies for Immunohistochemistry

| Genotype | Primary Solution | Secondary Solution |
| --- | --- | --- |
| *Opn4-GFP* (post-recording pieces) | Streptavidin 488 (Thermo, S11223), mouse anti-SMI32 (BioLegend, 801701), goat anti-ChAT (Milipore, AB144P) | Streptavidin 488, donkey anti-mouse (Thermo, A31571), donkey anti-goat (Thermo, A-11056) |
| *Opn4<sup>Cre/+</sup>;ZEG* | rabbit anti-GFP (Thermo, A11122), mouse anti-SMI32 | goat anti-rabbit (Thermo, A11034), goat anti-mouse (Thermo, A21125) |
| *Opn4<sup>LacZ/+</sup>;Opn4-GFP* | chicken anti-Beta galactosidase (Invitrogen, A11132), rabbit anti-GFP | goat anti-chicken (Thermo, SA172000), goat anti-rabbit (Thermo, A-11035) |
| *Opn4<sup>LacZ/+</sup> & Opn4<sup>Cre/+</sup>* | Mouse anti-Brn3a (Milipore, MAB1585), goat anti-ChAT | Donkey anti-mouse, donkey anti-goat |

All primary and secondary solutions are 0.3% Triton-X and 6% goat or donkey serum. With the exception of ChAT, all primary and matching secondary were done at 1:500 dilutions. ChAT and corresponding secondary were done at 1:250.

## Results

### Morphological properties of ipRGC subtypes during development

In order to assess the morphological and physiological properties of ipRGC subtypes during development, we first needed to confirm that we could reliably identify each subtype at early postnatal stages using criteria available to differentiate the adult subtypes. We chose to focus on M1, M2, and M4 ipRGCs because the properties of these subtypes are well characterized and they have been previously shown to tile the retina (25–27). M1, M2, and M4 ipRGCs can be differentiated by their dendritic stratification in the inner plexiform layer (M1: OFF stratifying and M2, M4: ON stratifying) and by presence (M4) or absence (M1, M2) of SMI-32 immunolabeling. We therefore first wanted to determine whether we could identify ipRGC subtypes during postnatal development using these same criteria: M1 ipRGCs, OFF stratifying and SMI-32 negative, M2 ipRGCs, ON stratifying, SMI-32 negative, and M4 ipRGCs, ON stratifying, SMI-32 positive. We targeted ipRGCs in Opn4-GFP mice for patch clamp recordings of ipRGCs at P6, P8, P10, P14, and Adult ages and filled cells with neurobiotin. We then performed immunohistochemistry for SMI-32 and choline acetyltransferase (ChAT), determined whether each cell was SMI-32 positive and whether it was ON or OFF stratifying (using ChAT bands as a reference). Using the aforementioned subtyping criteria, we find that we can indeed clearly identify these three ipRGC subtypes in our earliest time point, postnatal P6 (Figure 1). Interestingly, when we mapped the lamination patterns of M1, M2, and M4 ipRGCs at P6 and adulthood, we found that all ipRGC subtypes had a different lamination pattern compared to their adult counterparts with the M1 subtype being most similar to adulthood (Figure 1B). In contrast, the M2 and M4 subtypes seem to experience a bigger change in lamination pattern as cells mature. We observed that the M2 ipRGCs stratify closer to the middle of the IPL in early postnatal development before refining this dendritic lamination to the innermost portion of the IPL in adulthood (Figure 1D) and that the M4 ipRGCs have dendrites stratifying closer to the ganglion cell layer in early postnatal development but then moving slightly closer to the middle of the IPL in adulthood (Figure 1F), in agreement with previous observations of adult M2 and M4 ipRGC morphology (28). These findings suggest that although the M1, M2, and M4 ipRGCs’ dendrites broadly stratify within the correct layer early on, their dendritic stratification undergoes refinement in later parts of postnatal development.

**Figure 1:**
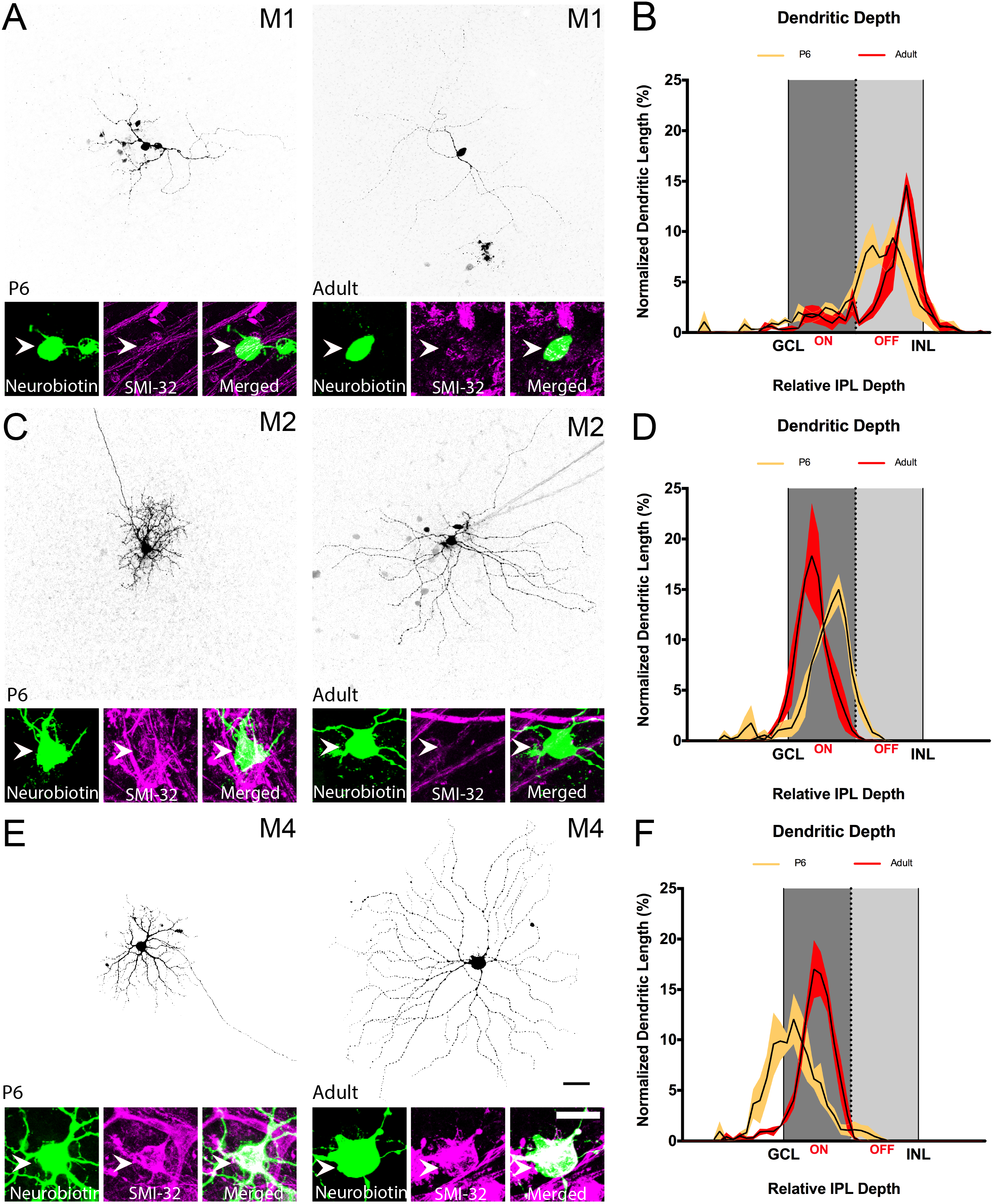
M1, M2, and M4 subtypes can be identified using immunohistochemistry and dendritic stratification at P6. (A) M1 ipRGCs filled with neurobiotin (top panel) at P6 (left) and Adult (right) stages. Cells were immunolabeled for SMI-32 (bottom panels). M1 ipRGCs are SMI-32 negative. (B) Dendritic depth measurements for M1 ipRGCs at P6 (yellow) and Adult (red) ages (n = 3 cells/age). (C) M2 ipRGCs filled with neurobiotin (top panel) at P6 (left) and Adult (right) stages. Cells were immunolabeled for SMI-32 (bottom panels). M2 ipRGCs are SMI-32 negative. (D) Dendritic depth measurements for M2 ipRGCs at P6 (yellow) and Adult (red) ages (n = 3 cells/age). (E) M4 ipRGCs filled with neurobiotin (top panel) at P6 (left) and Adult (right) stages. Cells were immunolabeled for SMI-32 (bottom panels). M4 ipRGCs are SMI-32 positive. (F) Dendritic depth measurements for M4 ipRGCs at P6 (yellow) and Adult (red) ages (n = 3 cells/ age). White arrows point to soma. Dark gray shading indicates ON sublamina, light gray indicates OFF. GCL and IPL refer to middle of respective cell body layer. Scale bar is 50µm.

The ability to define ipRGC subtypes early in development affords us the opportunity to characterize the progression of ipRGC structural and functional development in a way that is not possible for most RGC types. We first analyzed the morphological changes that occur in each ipRGC subtype during postnatal development. To do this, we filled M1, M2, and M4 ipRGCs with Neurobiotin at P6, 8, 10, 14 and Adult stages. We measured soma size, dendritic field diameter, and total dendritic length, and performed Sholl analysis to assess the complexity of the dendritic arbors (Figure 2). We found that soma size remained constant across development in M1 and M2 ipRGCs, but increased in M4 ipRGCs (Figure 2H). With regards to dendritic field size, we found that M1 ipRGCs exhibit adult dendritic field size and length by P10 (Figure 2C-D), while M2 ipRGCs mature by P14 (Figure 2F-G) and M4 ipRGCs continuing to expand their dendritic field size and complexity beyond P14 (Figure 2I-J).

**Figure 2:**
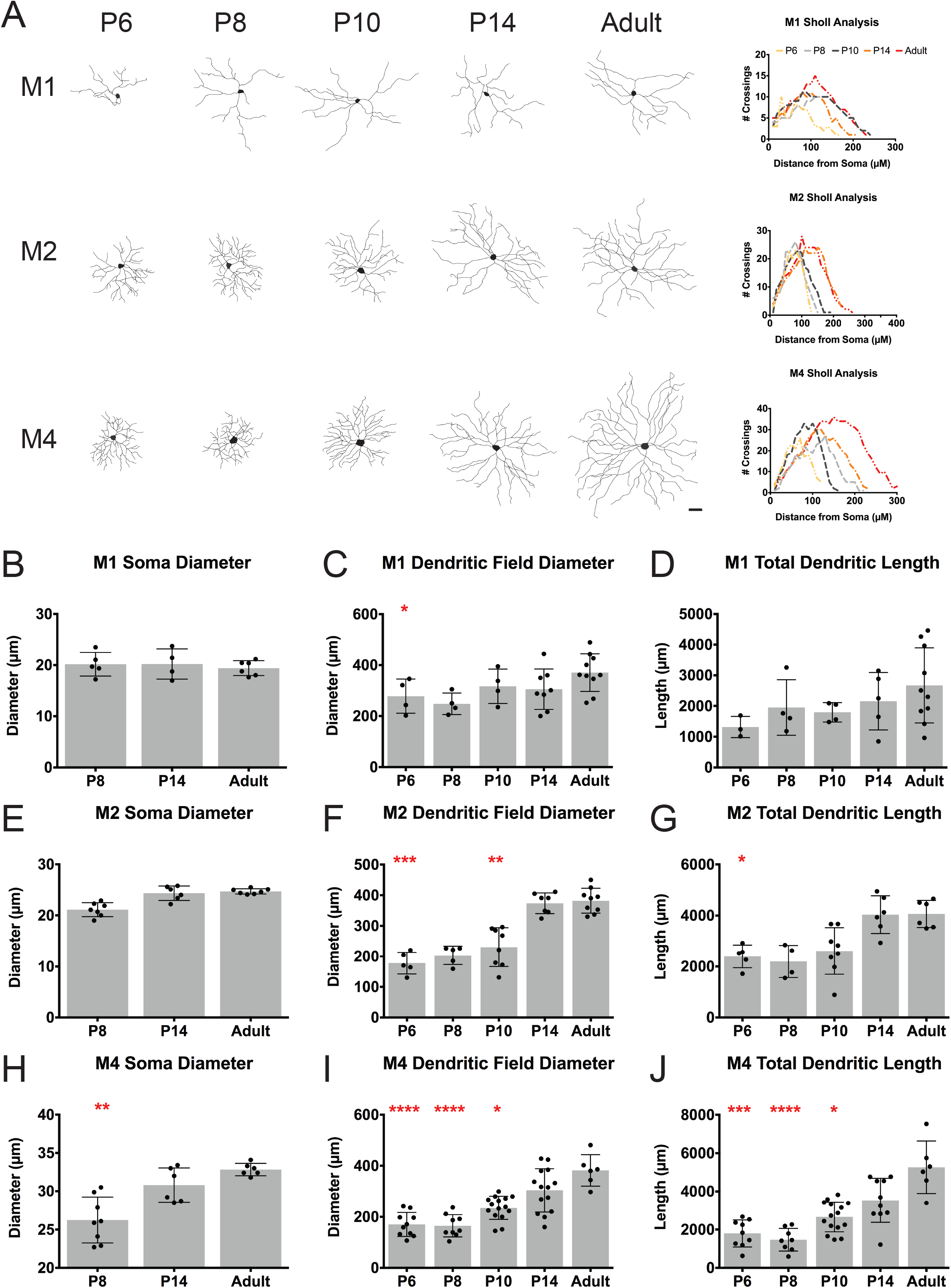
Dendritic length and diameter measurements for M1, M2, M4 subtypes during development. (A) Representative cell tracings of M1, M2, and M4 subtypes for P6, P8, P10, P14, Adult timepoints (*left*) and corresponding Sholl analysis for representative cells (*right*). (B-D) Mean ± SD M1 ipRGC soma diameter (B), dendritic field diameter (C), and total dendritic length (D). (E-G) Mean ± SD M2 ipRGC soma diameter (E), dendritic field diameter (F), and total dendritic length (G). (H-J) Mean ± SD M4 ipRGC soma diameter (H), dendritic field diameter (I), and total dendritic length (J). Scale bar is 50µm, n=5-8 per subtype/age,*p<0.05, **p<0.01, ***p<0.001, ****p<0.0001 when compared to Adult time point.

In adulthood, M1 ipRGCs have the smallest somata and smallest, least complex dendritic arbors amongst these three subtypes while M4 ipRGCs have the largest somata, as well as the largest and most complex dendritic arbors (1, 25, 28) (Figure 3F). We therefore next examined whether the reported morphological differences between adult ipRGC subtypes could be detected at early postnatal stages (Figure 3). Interestingly, at P8, we find that M1 ipRGCs have the largest dendritic field diameter while M4 ipRGCs have the smallest, which may be reflective of a faster rate of maturation for M1 ipRGC morphology (Figure 3A). All three subtypes exhibit similar total dendritic length at this age while in adulthood M4 cells have the largest total dendritic length of the three subtypes (Figure 3C-D). Of note, we found a large spread in the morphological measurements for the M1 and M4 subtypes the adult stage (Figure 4), and so we did not find that the subtypes were significantly different in dendritic field diameter (Figure 3B), as had been previously reported (1, 25, 28). The M4 variation is likely a function of the large differences in M4 ipRGC arbors from nasal, where M4 cells are very large, to temporal retina where M4 cells are very small (29). Additionally, M1 ipRGCs have been reported to show large variation in their morphological (and biophysical) properties (30). Sholl analyses comparing morphological complexity between all three subtypes reveals that the M2 and M4 ipRGCs begin to exhibit more complex dendritic arbors than M1 ipRGCs at early postnatal stages (Figures 3E-F).

**Figure 3:**
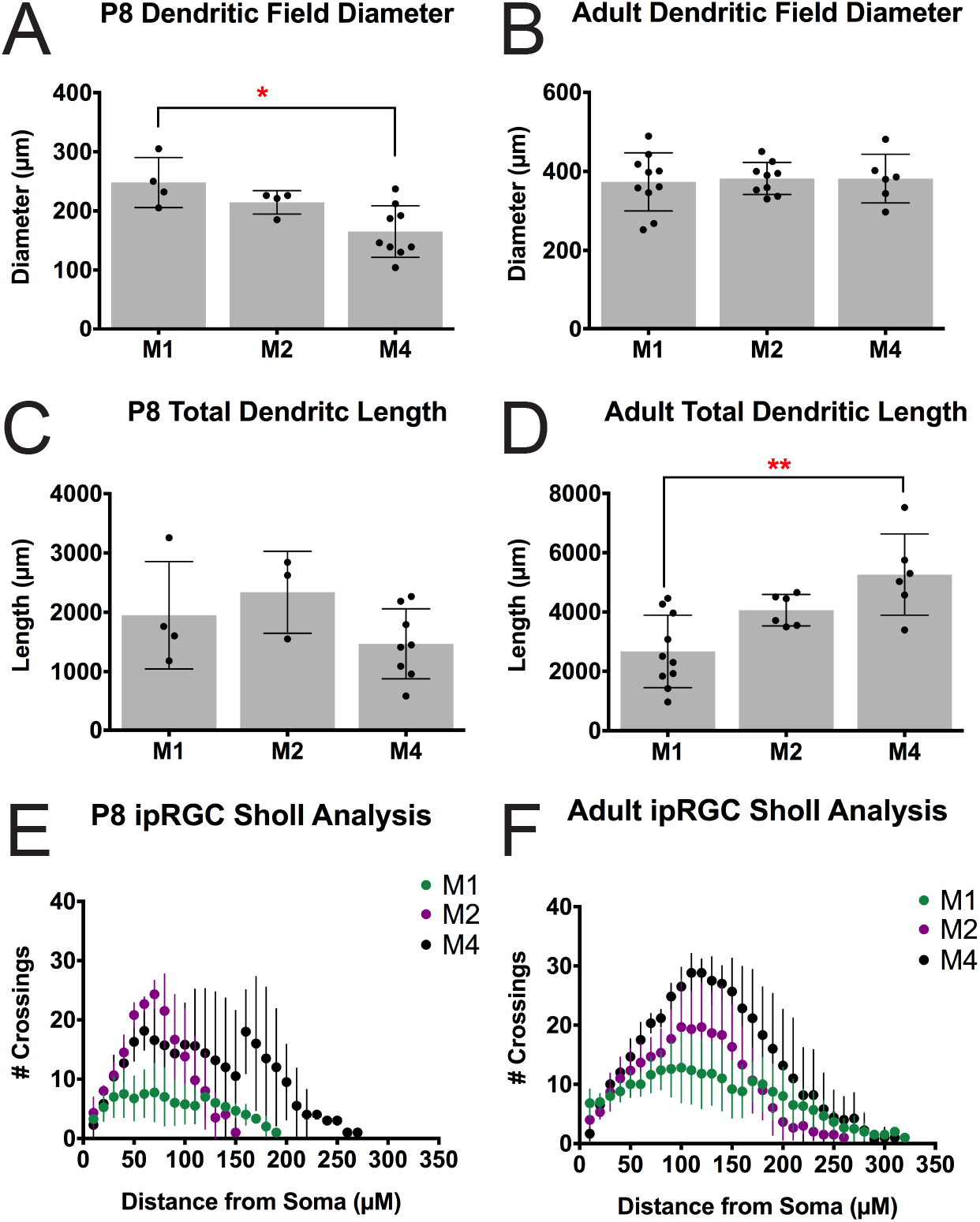
Comparison of ipRGC subtype morphology at P8 and in Adulthood. (A-B) Mean ± SD dendritic field diameter at P8 (A) and Adult (B) for M1, M2, and M4 ipRGCs. (C-D) Mean ± SD total dendritic length at P8 (C) and Adult (D) for M1, M2, and M4 ipRGCs. (E-F) Mean ± SD number of crossings in Sholl analysis of M1, M2, and M4 ipRGCs at P8 (E) and Adult (F). n=5-8 per subtype/age, *p<0.05, **p<0.01.

**Figure 4:**
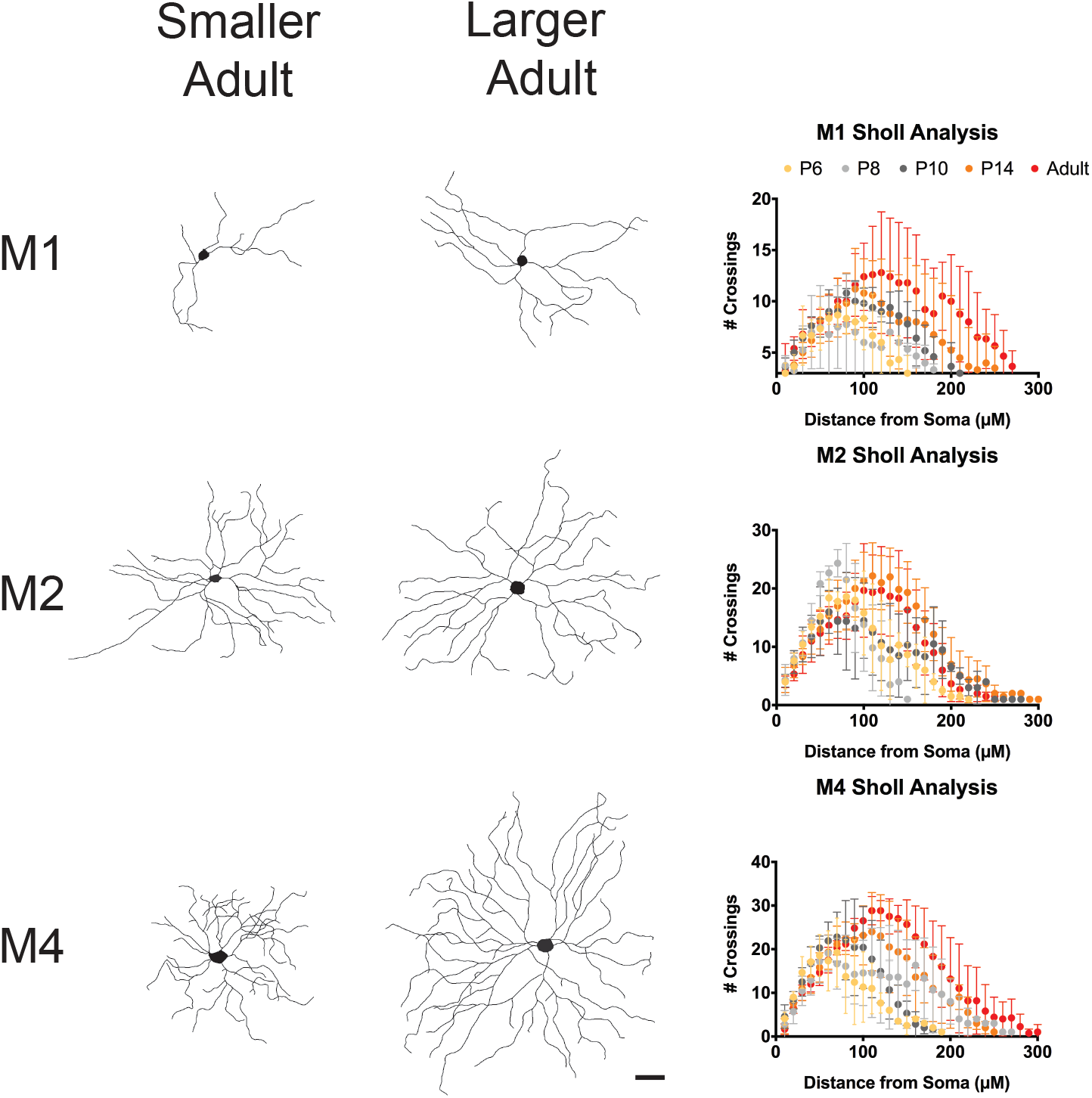
M1 and M4 subtypes have large variation of morphology in adulthood. *Left*, Representative cell tracings of small and large Adult cells for the M1, M2, and M4 subtypes. *Right*, Average Sholl analysis for M1, M2, and M4 subtypes at P6, P8, P10, P14, and Adult. Graphs are Mean ± SD, n=5-8 per subtype/ age. Scalebar is 50µm.

### Physiological properties of ipRGC subtypes during development

Following morphological analysis, we next characterized the intrinsic physiological properties of M1, M2, and M4 ipRGCs across development. In general, the intrinsic physiological properties of each subtype were relatively stable across development (Figure 5). We observed that M1 cells have a downward trend in capacitance and input resistance as cells age (Figure 5C-D) while M2 and M4 cells experience a drop in capacitance between P14 and adult, as well as a downward trend in input resistance as development progresses (Figure 5G-H, K-L). The variation in capacitance and resistance in particular are likely to be a combination of changes in membrane surface area, intrinsic membrane properties, and electrical coupling with a surrounding network of cells (31). When we directly compared the input resistance and resting membrane potential of M1, M2, and M4 ipRGC subtypes at P8 and Adult ages, we found that M1 cells have a more depolarized resting membrane potential and higher input resistance even early in development (Figure 6A, C). These differences mimic those previously observed in light adapted tissue for adult M1 versus M2 and M4 ipRGCs (20, 32) as well as our own observations (Figure 6B, D).

**Figure 5:**
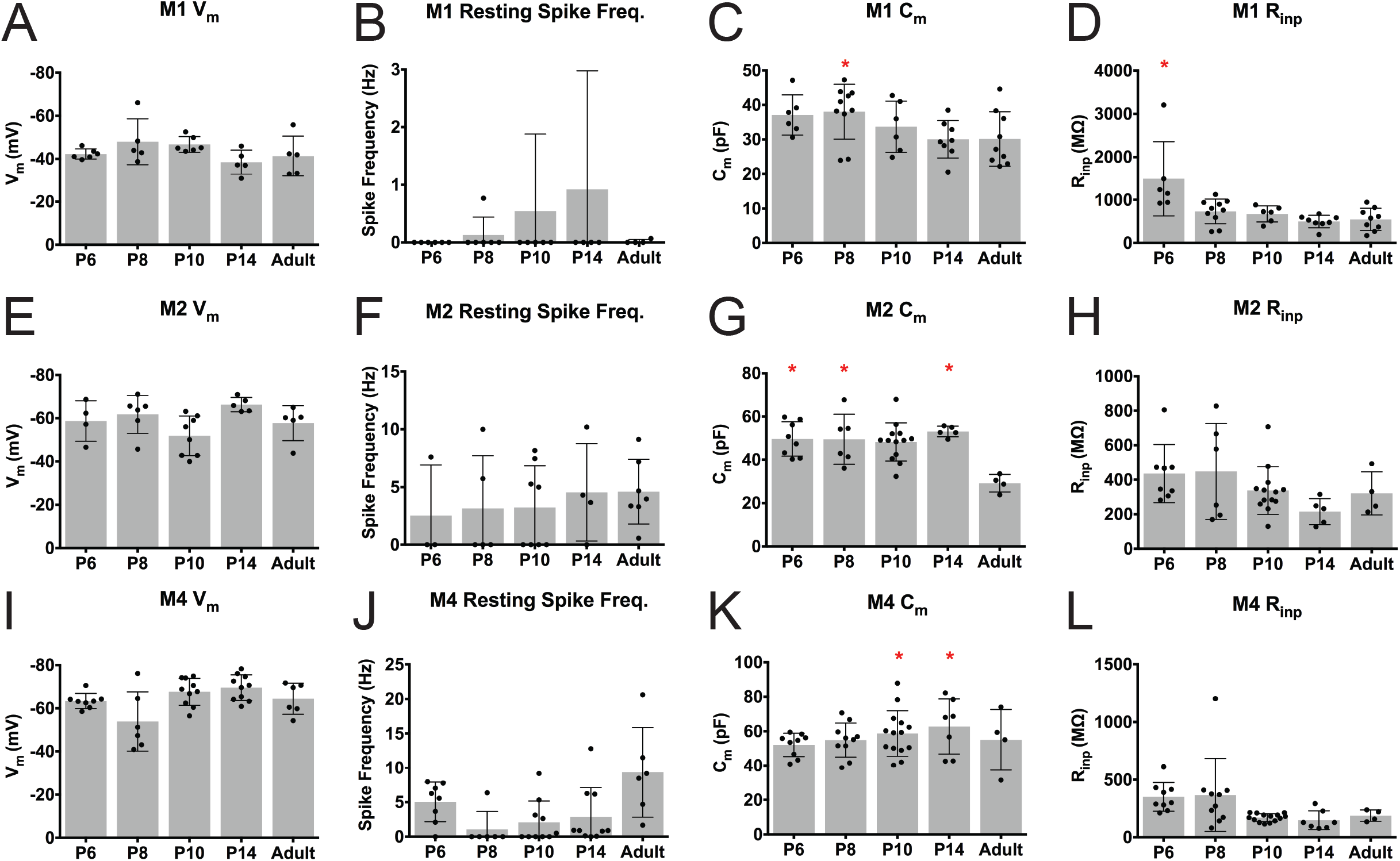
Intrinsic physiological properties of ipRGC subtypes across development. (A-D) Mean ± SD M1 ipRGC resting membrane potential (A), spike frequency at rest (B), capacitance (C), and input resistance (D). (E-H) Mean ± SD M2 ipRGC resting membrane potential (E), spike frequency at rest (F), capacitance (G), and input resistance (H). (I-L) Mean ± SD M4 ipRGC resting membrane potential (I), spike frequency at rest (J), capacitance (K), and input resistance (L). n= 5-14 per subtype/age *p<0.05 when compared to Adult time point.

**Figure 6:**
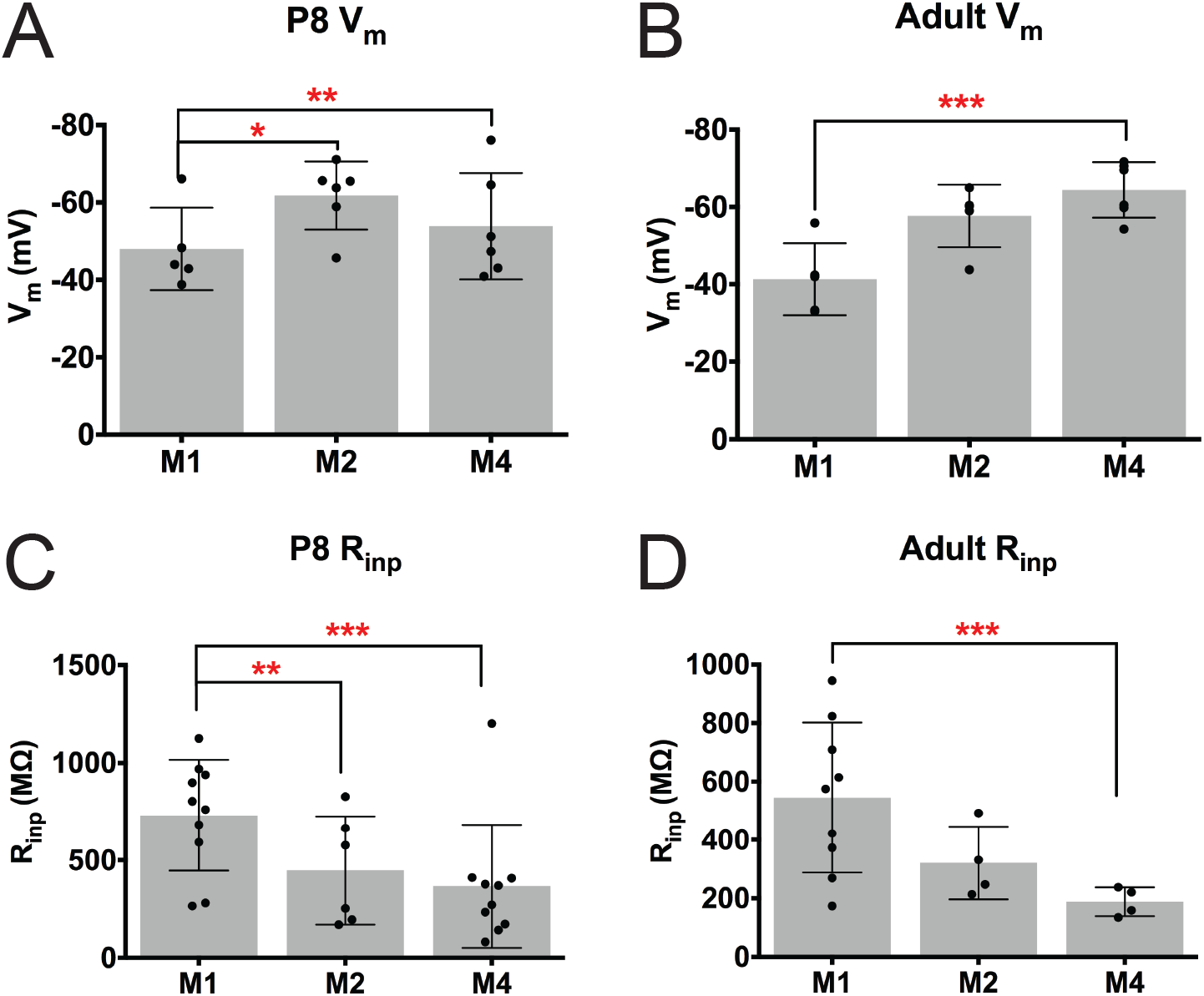
ipRGC subtypes exhibit distinct intrinsic properties from early postnatal development. (A-B) Mean ± SD resting membrane potential at P8 (A) and Adult (B) for M1, M2, and M4 ipRGCs. (C-D) Mean ± SD input resistance at P8 (C) and Adult (D) for M1, M2, and M4 ipRGCs. n= 5-14 per subtype/age, *p<0.05, **p<0.01, ***p<0.001.

We next compared the spiking properties and action potential waveform of M1, M2, and M4 ipRGCs. We performed current clamp recordings from each of these subtypes and injected 1s stepwise depolarizing current of 10 or 20pA until cells reached depolarization block. M1 ipRGCs show very few action potentials evoked by positive current (Figure 7A-B), as reported previously for light-adapted M1 cells (9, 20). In contrast, the M2 and M4 subtypes are much more excitable during development with the M4 subtype significantly increasing in excitability as cells mature (Figure 7A, D, F). Somewhat surprisingly, the current density needed to reach the maximum spiking frequency was not significantly different across ages (Figure 7C, E, G) for each of the subtypes. We next analyzed several components of individual action potentials from each subtype including width at half max, threshold, and fast after hyperpolarization (Figure 8). Unsurprisingly, we find that action potential width at half-max decreases for all cell types across development (Figure 8B, E, H) which is in line with typical progression of neuronal development (33, 34). We also observe that threshold decreases for the M2 and M4 subtypes as cells mature (Figure 8F, I).

**Figure 7:**
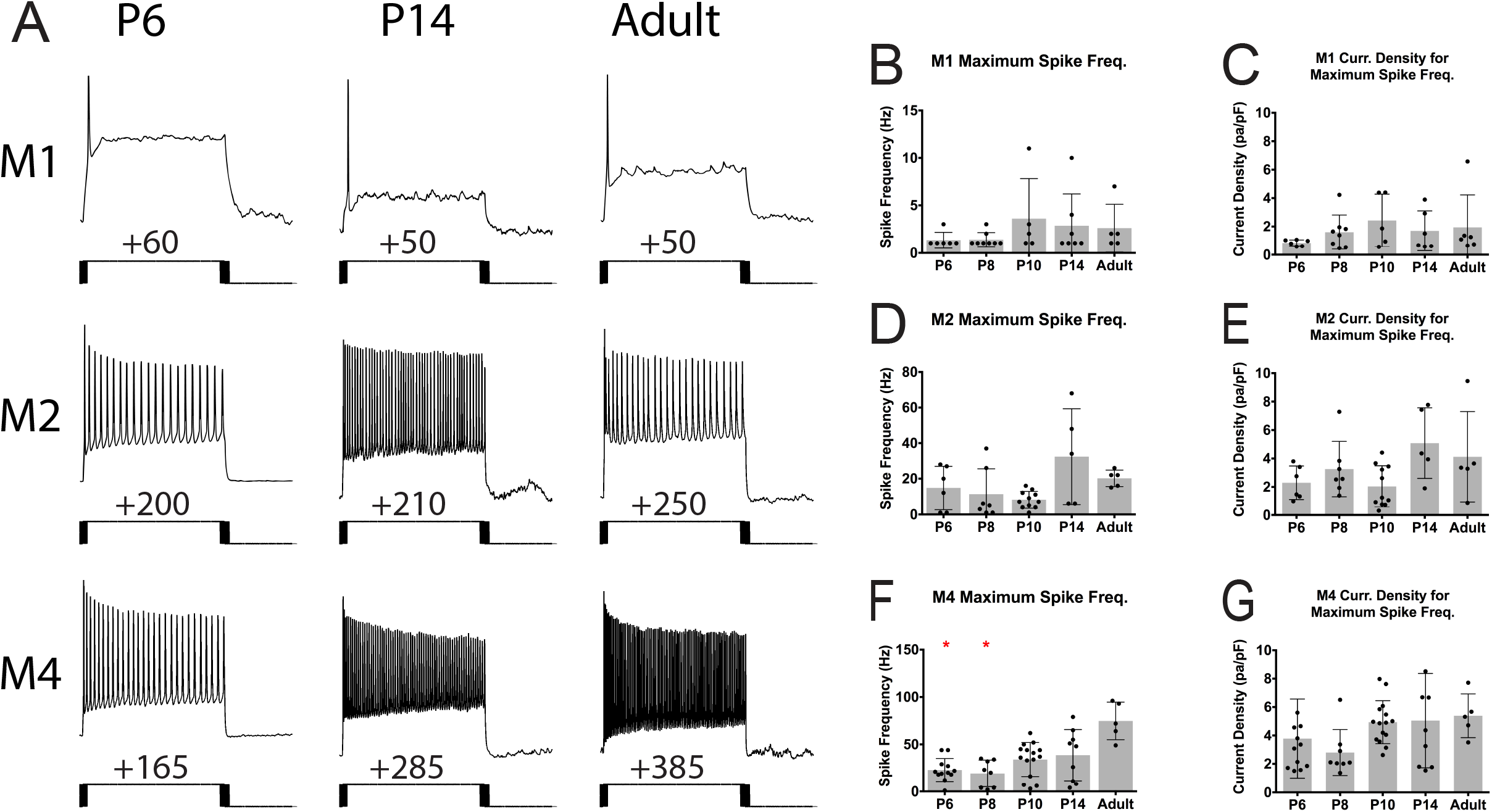
ipRGC subtype excitability across development. (A) Representative traces from depolarizing current injection that elicited the maximum spike output for M1, M2, and M4 subtypes at P6, P14, and Adult timepoints. (B-C) Maximum spike frequency elicited by depolarizing current steps (B) and maximum current density required to elicit maximum firing frequency (C) in M1 ipRGCs. (D-E) Maximum spike frequency elicited by depolarizing current steps (D) and maximum current density required to elicit maximum firing frequency (E) in M2 ipRGCs. (F-G) Maximum spike frequency elicited by depolarizing current steps (F) and maximum current density required to elicit maximum firing frequency (G) in M4 ipRGCs. Graphs are Mean ± SD, n=5-14 per subtype/age, *p<0.05 when compared to Adult timepoint.

**Figure 8:**
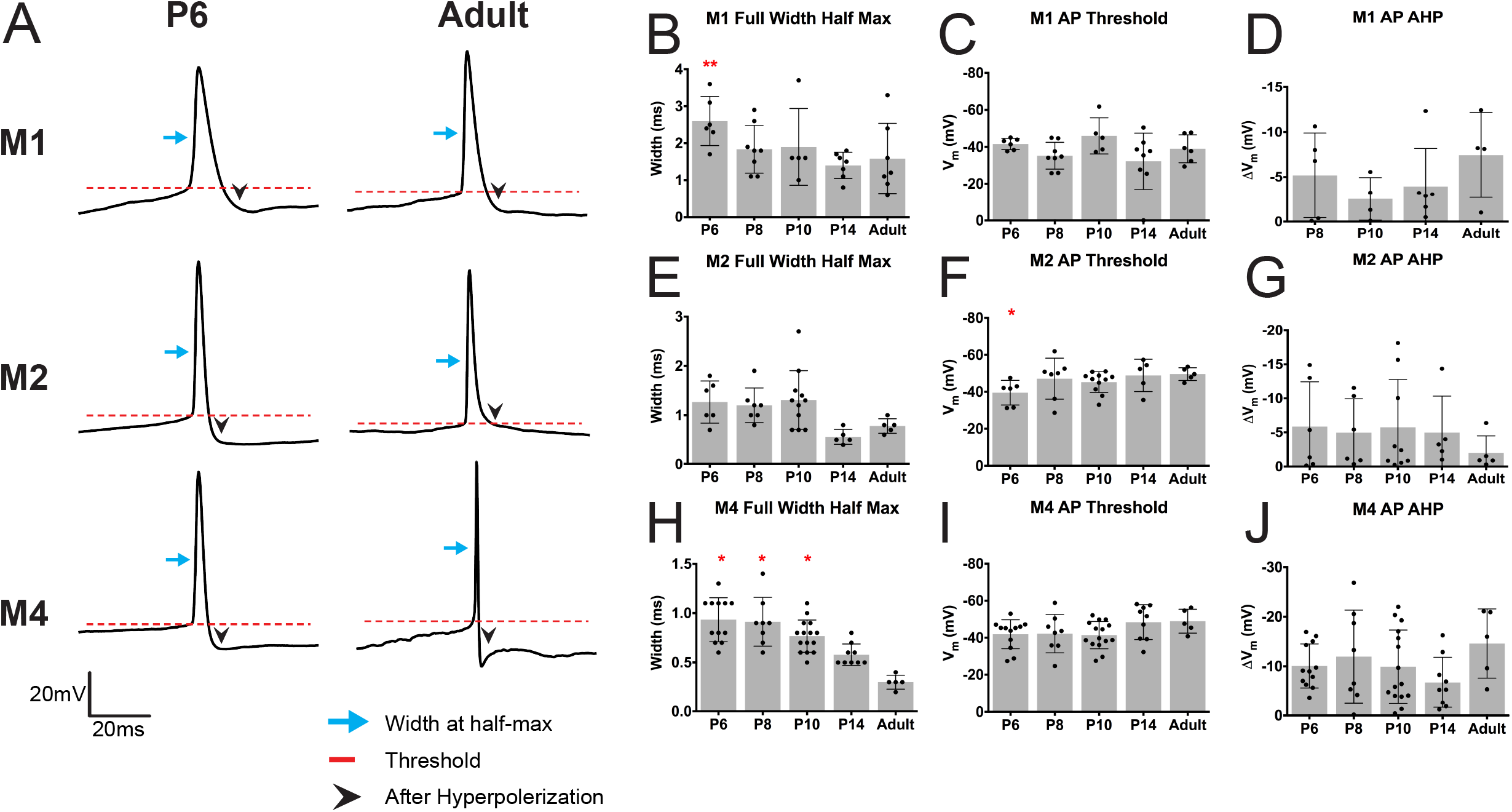
Action potential properties of ipRGC subtypes across development. (A) Representative traces of single action potentials for M1, M2, and M4 subtypes for P6 and Adult timepoints. (B-D) Measurement of M1 ipRGC AP full width at half max (B), AP threshold (C), and AP AHP (D). AP AHP P6 for M1 subtype is not plotted because cells did not hyperpolarize after action potential elicitation. (E-G) Measurement of M2 ipRGC AP full width at half max (E), AP threshold (F), and AP AHP (G). (H-J) Measurement of M4 ipRGC AP full width at half max (H), AP threshold (I), and AP AHP (J). Graphs are Mean ± SD, n=5-14 per subtype/age, *p<0.05 when compared to Adult timepoint. AP: action potential, AHP: after hyperpolarization. Blue arrow indicates width at half-max, red dotted line indicates threshold, and black arrowhead indicates after hyperpolarization.

In addition to the intrinsic properties of M1, M2, and M4 ipRGCs, we also examined the ipRGC light response across development. We performed current clamp recordings of ipRGC light responses to 30s of saturating blue light stimulus at 1×10^17^ photons/cm^2^s^−1^ at P6, P8, P10, P14, and Adult. In general, we found that all subtypes exhibited adult-like light responses by P14 (Figure 9), consistent with the intact synaptic circuitry in the retina around the time of eye opening(15, 16). Specifically, M2 and M4 ipRGCs, which are known to receive strong drive from the cone pathway (1, 28, 35), show faster and larger light responses as development progress (Figure 9D-G). M1 ipRGCs, however, had statistically similar light responses throughout development (Figure 9B-C). This is in line with previous reports that M1 ipRGCs are strongly driven by melanopsin phototransduction in bright light (35), and indicates that M1 ipRGCs show mature light responses from early developmental stages. M1 cells also showed strong depolarization block in their light responses, as reported previously (36). Interestingly, when we compared ipRGC light responses early in development and adulthood, we observe that while the maximum depolarization in response to light is similar between all three subtypes in both adulthood and development (Figure 10A-B), the M1 subtype has a faster onset time early in development, but responds more slowly than other subtypes in adulthood (Figure 10C-D).

**Figure 9:**
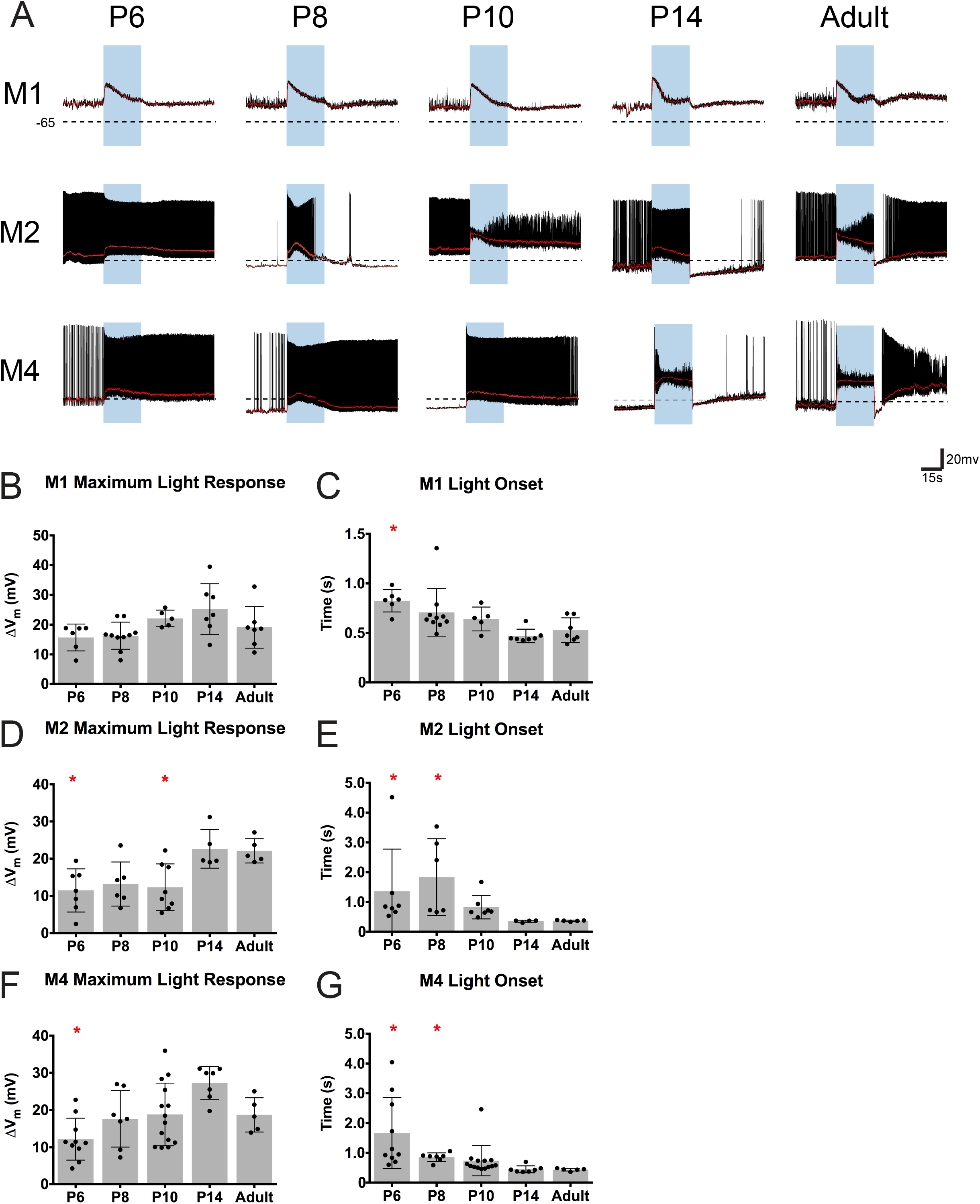
ipRGCs light responses across development. (A) Representative light response traces from M1, M2, and M4 cells at P6, P8, P10, P14, and Adult timepoints. Blue rectangle indicates start and end of light stimulus. Black dotted line indicates −65 mV. (B-C) M1 ipRGC maximum light response (B) and light onset (C). (D-E) M2 ipRGC maximum light response (D) and light onset (E). (F-G) M4 ipRGC maximum light response (F) and light onset (G). Light onset was defined by time to reach 50% of maximum depolarization. Graphs are Mean ± SD, n=5-14 per subtype/age, *p<0.05, **p<0.01 when compared to Adult time point.

**Figure 10:**
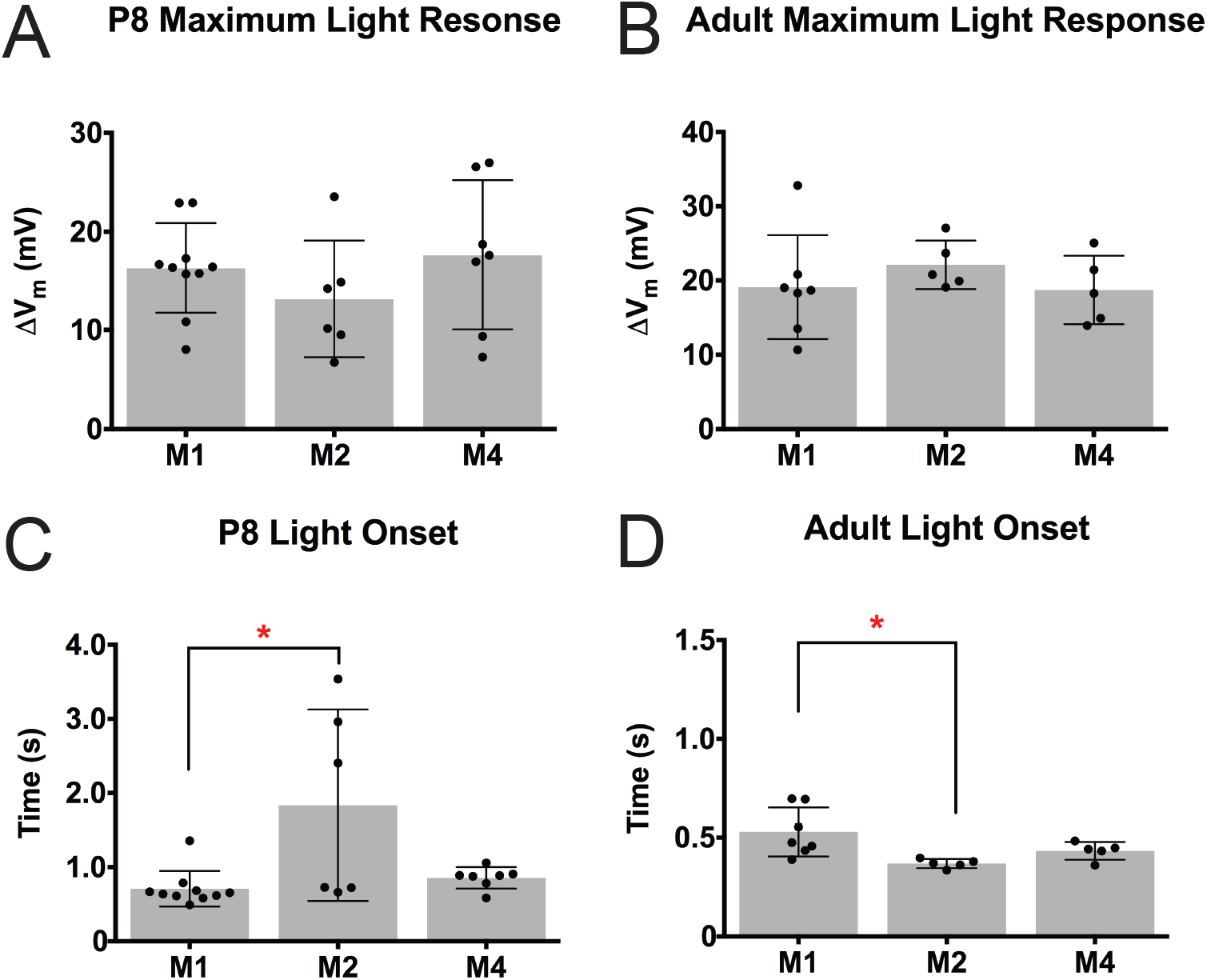
Comparison of light response properties of ipRGC subtypes across development. (A-B) Maximum light response of M1, M2, and M4 ipRGC subtypes at P8 (A) and Adult (B). (C-D) Light onset for M1, M2, and M4 ipRGC subtypes at P8 (C) and Adult (D). Light onset was defined by time to reach 50% of maximum depolarization. Graphs are Mean ± SD, n=5-14 per subtype/age, *p<0.05.

### Assessing the embryonic birthdate of ipRGC subtypes

Overall, our results suggest that ipRGC subtypes mature at different rates during postnatal development. We next asked whether these differences in maturation rate might be reflected in differences in cellular birthdate. That is, do the M1, M2, and M4 subtypes terminally differentiate at different embryonic timepoints, and how does this compare to the birthdate of conventional RGCs? To answer this, we utilized 5-ethynyl-2’-deoxyuridine (EdU), a thymidine analog, to label cells that terminally differentiated on specific embryonic days, also known as birthdating. We first compared the birthdate of all ipRGCs, M1-3 ipRGCs, and Brn3a-positive RGCs (non-ipRGCs). To do this, we quantified the percentage of cells that were EdU and GFP positive in both *Opn4^Cre/+^; Z/EG* animals (where all ipRGCs are labeled with GFP; Figure 11A) and *Opn4^LacZ/+^; Opn4-GFP* animals (where only M1-M3 ipRGCs are labeled with GFP and only M1 ipRGCs are labeled with LacZ; Figure 11A) from Embryonic Day E11-14. We also immunostained for a non-ipRGC population of RGC, the Brn3a-positive RGCs (Figure 11A), and counted the number EdU-positive, Brn3a-positive RGCs from E11-14. While ipRGCs appear to be born primarily on E11 and E12 (Figure 11B-C), we observed that Brn3a positive RGCs continued to terminally differentiate at E13 and E14, suggesting that ipRGC birthdates differ from other RGC types.

**Figure 11:**
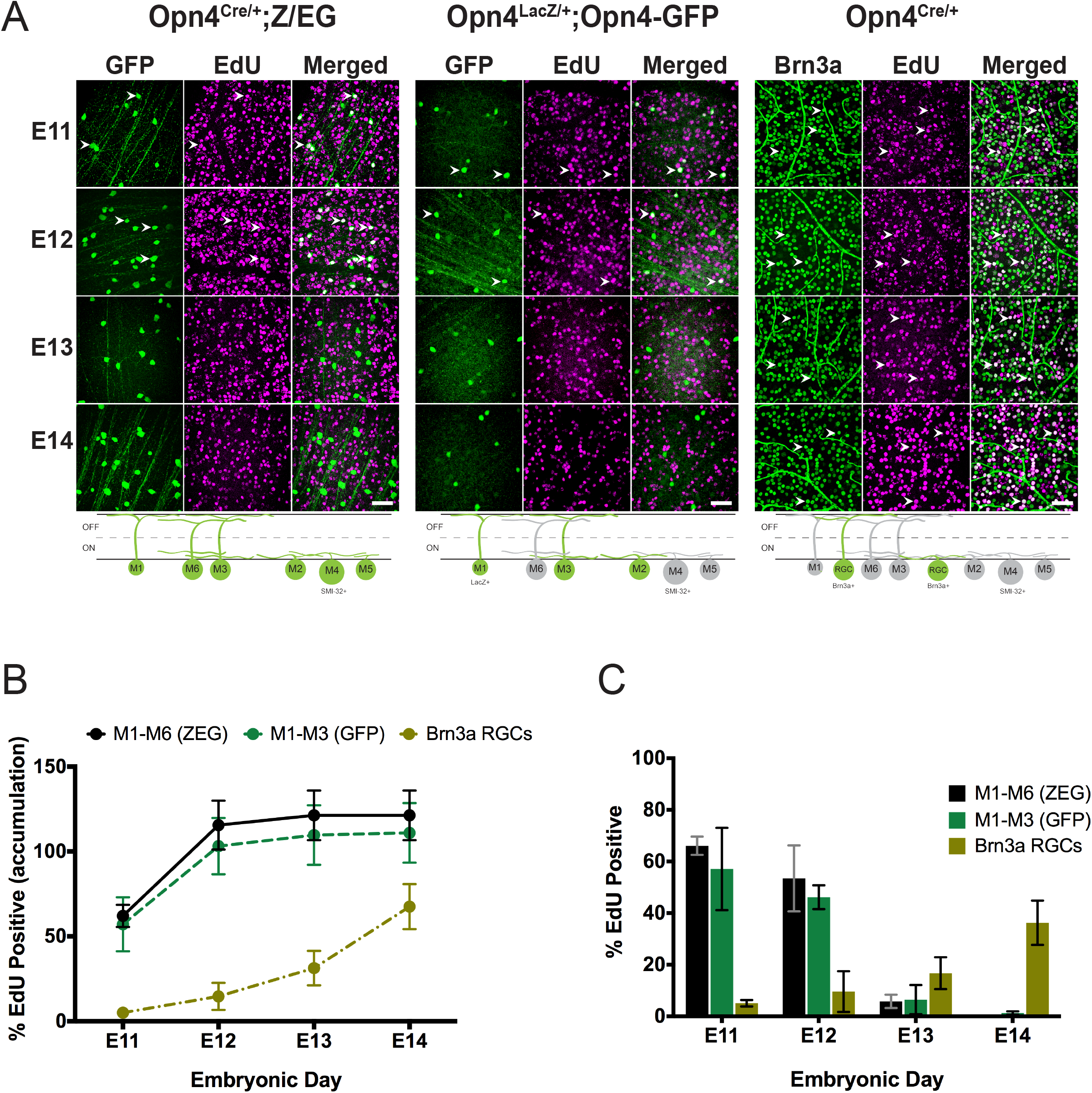
ipRGCs are born earlier than Brn3a positive RGCs. (A), *Top*, GFP or Brn3a immunohistochemistry (green) of retinas from Adult *Opn4*^*Cre*/+^;*ZEG*, *Opn4*^*LacZ*/+^; *Opn4-GFP* and *Opn4*^*Cre*/+^ animals exposed to EdU (magenta) at different developmental stages. White arrow heads point to examples cells co-labeled with GFP or Brn3a and EdU. *Bottom*, Schematic of cell types labeled green in each experiment. (B) Accumulation plot of proportion of ipRGCs or Brn3a RGCs that are EdU positive when exposed to EdU at different embryonic timepoints. Accumulation was calculated based on adding together the proportion data calculated for each timepoint. (C) Proportion of ipRGCs or Brn3a RGCs that are EdU positive when exposed to EdU on specific embryonic day. Graphs are Mean ± SD, n=3-4 retinas per timepoint. Scale bar is 100µm.

We next wanted to assess and compare the birthdate of individual ipRGC subtypes (M1, M2/3, and M4 ipRGCs). To identify M4 ipRGCs from “non-M4” ipRGCs, we immunolabeled *Opn4^Cre/+^; Z/EG* retinas for SMI-32. M4 ipRGCs are easily identified as GFP positive, SMI-32 positive, while non-M4 ipRGCs are GFP positive, SMI-32 negative (Figure 12B). In this line, OFF alpha RGCs can also be identified as GFP negative, SMI-32 positive. To differentiate M1 and M2/3 ipRGCs, we immunolabeled *Opn4^LacZ/+^; Opn4-GFP* mice for LacZ and GFP. M1 ipRGCs will be both GFP and LacZ positive (Figure 12A), while M2/3 ipRGCs should be GFP positive, LacZ negative (though some M3 ipRGCs may be LacZ positive, see (37); Figure 12A). In agreement with our broad comparisons in Figure 11, we find that M1, M2, and M4 ipRGCs are all primarily born on E11 and E12 (Figure 12C-D). Interestingly, when we compared the birthdate of M4/ON alpha RGCs and OFF alpha RGCs, we find that the OFF alpha RGCs continue to be born through E13 (Figure 12E-F), highlighting an important difference in birthdate between the ON and OFF alpha RGC population, despite these cells being considered part of the same class of (alpha) RGC.

**Figure 12:**
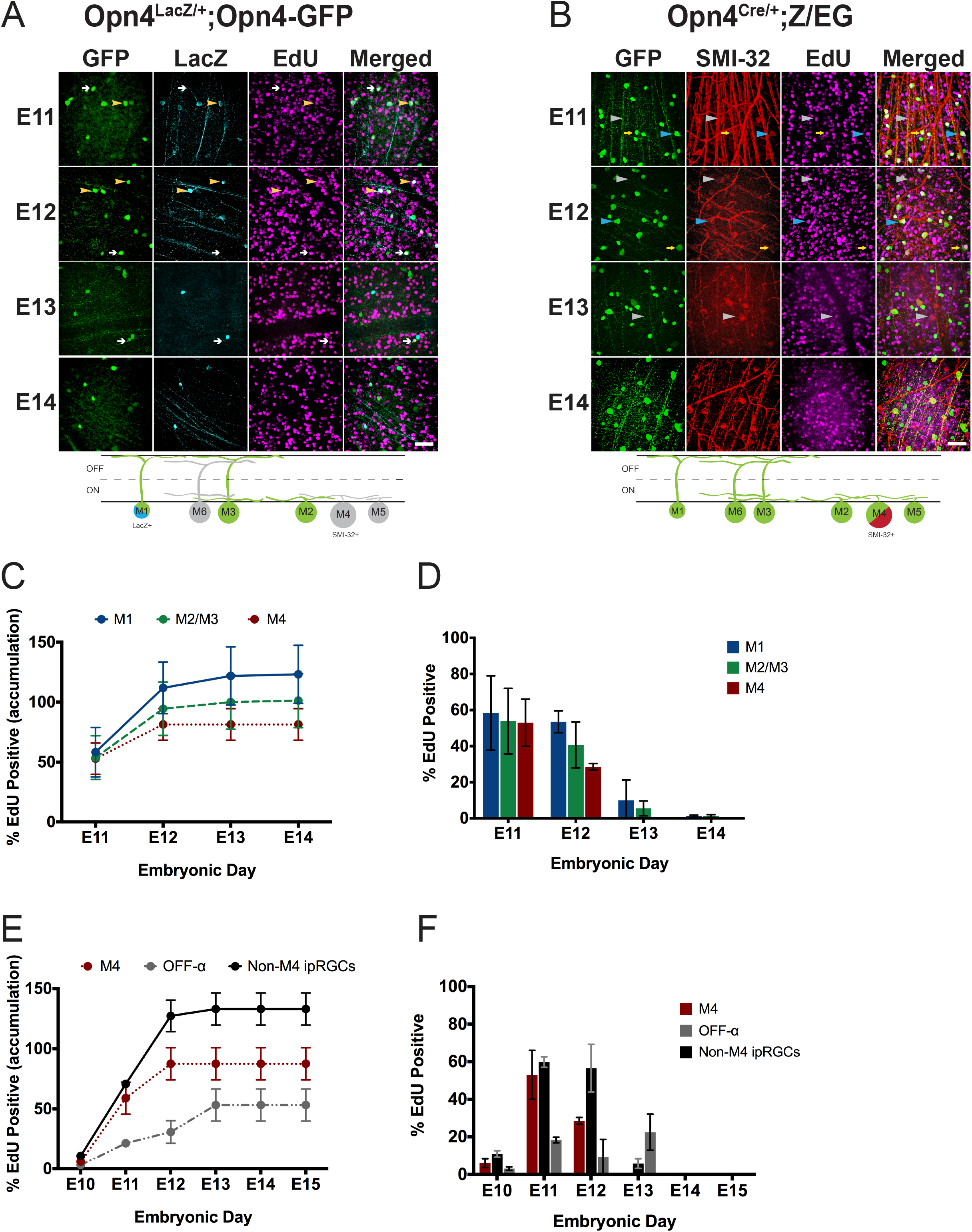
ipRGC subtypes are born at the same rate and frequency. (A) *Top*, GFP (green) and LacZ (cyan) immunohistochemistry in Adult *Opn4*^*LacZ*/+^; *Opn4-GFP* retinas labeled for EdU (magenta). Yellow arrowheads point to EdU positive M1 cells (GFP+, LacZ+) and white arrows to EdU-positive M2 cells (GFP+, LacZ−). *Bottom*, Schematic of ipRGC subtypes labeled with each marker in experiment. (B) GFP (green) and SMI-32 (red) immunohistochemistry in Adult *Opn4*^*Cre*/+^;ZEG retinas labeled for EdU (magenta). Blue arrowheads indicate EdU-positive M4 cells (GFP+, SMI-32+), yellow arrows indicate EdU positive non-M4 ipRGCs (GFP+, SMI-32−), and grey arrowheads indicate EdU positive OFF-alpha RGCs (GFP−, SMI-32+). *Bottom*, Schematic of ipRGC subtypes labeled with each marker in experiment. (C) Accumulation plot of proportion of ipRGC subtypes that are EdU positive when exposed to EdU at different embryonic timepoints. Accumulation was calculated based on adding together the proportion data calculated for each timepoint. (D) Proportion of ipRGC subtypes that are EdU positive when exposed to EdU at specific embryonic timepoints. (E) Accumulation plot of proportion of M4 ipRGCs, non-M4 ipRGCs, and OFF alpha RGCs that are EdU positive when exposed to EdU at different embryonic timepoints. Accumulation was calculated based on adding together the proportion data calculated for each timepoint. (F) Proportion of M4 ipRGCs, non-M4 ipRGCs, and OFF alpha RGCs that are EdU positive when exposed to EdU at specific embryonic timepoints. Graphs are Mean ± SD, n=3-4 retinas per timepoint. Scale bar is 100µm.

## Discussion

ipRGCs are a diverse class of RGC that influence not only a wide range of visual behaviors, but also several important components of retinal development such as spontaneous retinal waves and pruning of retinal vasculature. While there have been several studies that have looked at ipRGCs during postnatal retinal development, only a few have distinguished amongst ipRGC subtypes. Understanding the properties of ipRGC subtypes throughout development, as we have done in this work, is a crucial first step in defining the mechanisms by which ipRGCs exert their many and varied influences on retinal and visual system development.

### M1 ipRGCs have the largest dendritic field during early postnatal development

In general, we observed that all subtypes have the same upward trend in dendritic field expansion and complexity across early postnatal development. Surprisingly, we found that in early development the M1 subtype had the largest dendritic field and M4 cells had the smallest, despite a reversal of these patterns in adulthood (25, 28). However, we do find that in early development, like in adulthood, M1 cells have the least complex dendritic field (Figure 3; (20, 25, 28, 38)). Overall, our data suggest that the M1 subtype reaches an adult morphology earlier than either M2 or M4 cells. This could be attributed to the fact that M1 dendrites most likely undergo less expansion and branching relative to the M2 and M4 subtypes.

### Physiological properties are largely stable from early developmental stages

Unlike with morphology, we see that most of the general rules for physiological differences between the subtypes in adult animals are also observed in early postnatal stages. For example, the adult M4 subtype has been shown to be more excitable than M1 and M2 cells (9, 20, 32) and here we report that the M4 subtype is the most excitable among the three subtypes in both adulthood and during postnatal development (Figure 7). We also find that, like in adulthood, the M1 subtype has the most depolarized resting membrane potential and largest input resistance among ipRGC subtypes during early postnatal development (Figure 6; (9, 20, 32)). While it is expected that physiological properties for ipRGC subtypes would be different from what has been reported in adulthood, it is interesting that the physiological differences between subtypes remains relatively consistent through development. These findings support the idea that different subtypes might be influencing different aspects of retinal development via unique signaling properties and physiological roles. One other interesting observation that we note is that both input resistance and capacitance decrease in all subtypes as cells mature, although it is more gradual in the M1 subtype relative the M2 and M4 subtypes (Figure 5). Changes to capacitance and, to some extent, input resistance are indicative of changes in amount of cellular membrane surface area. Given that we observe an overall growth of the dendritic field and thus an increase in membrane for all subtypes, we would expect capacitance to increase as cells mature. The fact that we observe the exact opposite of this indicates that membrane space must be decreasing in some other way that we did not observe morphologically. One such way would be changes in electrical coupling between cells, which can influence capacitance and input resistance. In fact, it has been reported that ipRGCs during development are extensively coupled (12, 31). While there has yet to be a study that directly looks at how coupling changes in ipRGC subtypes across development as well as how it differs between subtypes during development, it has been shown that M1 and M2 ipRGCs are coupled to GABAergic and ON displaced amacrine cells in adulthood (38, 39). Similarly, adult M4 cells have been shown to couple amacrine cells (40). In contrast, work done by Arroyo et al, revealed that during development, ipRGCs are mostly connected to other retinal ganglion cells and other ipRGCs with low connectivity to GABAergic and other types of amacrine cells (31). They also showed that the number of cells that ipRGCs couple to 15 cells on average. In comparison, ipRGCs in adulthood have been found to couple to 5-25 cells with stark differences in number of cells coupled between subtypes (38). Taken together, this suggests that there is most likely a profound change in coupling between development and adulthood, a phenomenon that has been reported for ON-OFF direction-selective RGCs (41). Further work will need to be done to understand how the network changes as development progresses and if it changes different from subtype to subtype.

### Diversity ipRGC light responses during development

Multielectrode array recordings of light responses in P8 retinas were one of the first ways in which it was revealed that there are multiple subtypes of ipRGCs. Tu et al found that there were three types during development based on light onset as defined by start of spike output: Type I, slow onset, sensitive, fast offset, Type II, slow onset, insensitive, slow offset, and Type III, rapid onset, sensitive, very slow offset (13). Follow up studies have suggested that adult M1 is type III (20) and adult M2 is type II (20) and adult M4 is type I (14). In complement to this, we used whole cell recording techniques to show that maximum depolarization is similar across subtypes at P8 and that when we define light onset by time to reach 50% of maximum light response, we find that M1 subtype (Type III) is still the fastest with the M2 (Type II) and M4 (Type I) subtypes having similar onset times (Figure 10). Combined, this illustrates that while subtypes have similar maximum depolarizations in response to light, the kinetics of that response are actually very different. This diversity in kinetics and firing frequency gives rise to the very likely possibility that different subtypes might be modulating different developmental factors in response to light. However, it is not clear which components of the light response (firing frequency, spike latency, onset time of maximum response, or absolute maximum depolarization) are important determinants in modulating different aspects of retinal development in response to light and if the determining feature varies between light responsive developmental traits. Currently, it seems to be that any or all of the ipRGC subtypes could be the modulators of retinal vasculature or the prolonging retinal waves in response to light. Genetic models that allow us to ablate single subtypes or abolish the melanopsin response within a particular subtype will help resolve the requirements of the melanopsin response as well as which subtypes are necessary for specific behaviors.

### ipRGC birthdates diverge from conventional RGCs

Previously, it had been reported that RGCs have different birthdates based on their ganglion cell classification (42) and that Cdh3 positive RGCs which include a subset of the ipRGC population (43) are born between E10 and E12. Furthermore, it has also been reported that the majority M1 ipRGCs are born between E11 and E12 (44). However, this study counted LacZ+ ipRGCs at P0, a time point at which other subtypes have been reported to express high amounts melanopsin (14). Thus, making it unclear if this was a purely M1 ipRGC population. Nonetheless, it is clear that some ipRGCs are born in the earlier part of retina cell type neurogenesis. Given the morphological and physiological differences within ipRGC subtypes (Figures 3, 6; (20, 25, 28)), we wondered whether non-M1 ipRGCs would also be born in the E11-E12 timeframe or if they would have different birthdates. Our results show that the majority of the M1, M2, and M4 ipRGC subtypes are born in within the E10-E12 timeframe and also reveal that M1, M2, and, M4 ipRGCs are all born at same rate (Figure 12). The study done by Osterhout in 2014 also showed that the time at which an RGC is born can dictate the strategy the cell will employ in axon targeting. ipRGC subtypes each target very different brain regions, with M1 and some M2 ipRGCs targeting non-image forming targets and other M2 and M4 ipRGCs targeting image-forming brain regions (25, 26, 45), indicating that RGCs with different downstream targets are also born at overlapping time points. Interestingly, we also observed that OFF alpha ganglion cells which share the alpha RGC classification with the M4 subtype, show distinctly different birthdating patterns from the ON alpha RGCs. It is possible that these temporal differences in differentiation underlie additional differences in ON versus OFF alpha RGC properties.

## Conclusions

Because most RGC types are identified based on their adult characteristics, following distinct RGC types across development has proven difficult. ipRGCs early expression of melanopsin along with other identification markers provide us a unique opportunity to follow multiple RGC subtypes through development. Leveraging this advantage, we were able to carry out a broad characterization of ipRGC subtypes throughout the developmental timepoints at which they are influencing retinal and visual system development. This study lays the groundwork for future studies into the precise role of each ipRGC subtype in retinal development.

## List of Abbreviations

ipRGC: intrinsically photosensitive retinal ganglion cells
RGC: retinal ganglion cells
ChAT: choline acetyltransferase
IPL: inner plexiform layer
GCL: ganglion cell layer
INL: inner nuclear layer

## Declarations

## Ethics and consent to participate

All procedures were approved by the Animal Care and Use Committee at Northwestern University.

## Consent for publication

N/A

## Availability of data and material

The datasets used and/or analyzed during the current study are available from the corresponding author on reasonable request.

## Authors’ contributions

JAL and TMS designed experiments, wrote paper, and prepared the figures. JAL collected and analyzed all of the data for the experiments.

## Acknowledgements

We would like to thank Dr. Gregory Schwartz and members of the Schmidt lab for helpful comments on the manuscript. We would also like to thank Dr. Marla Feller for the gift of the *Opn4-GFP* mice and Dr. Samer Hattar for the gift of the *Opn4*^*Cre*^ and *Opn4*^*LacZ*^ lines. We would also like to thank David Swygart for his help with the IPL depth analysis.

